# Estimating scale-dependent covariate responses using two-dimensional diffusion derived from the SPDE method

**DOI:** 10.1101/2024.12.17.628864

**Authors:** Max Lindmark, Sean C. Anderson, James T. Thorson

## Abstract

1. Species distribution models (SDMs) are widely used to standardize spatially unbalanced data, project climate impacts, and identify habitat for conservation. SDMs typically estimate the impact of local environmental conditions by estimating a dome-shaped or non-parametric “environmental response function”. However, ecological responses often integrate across local habitat conditions, such that species density depends on habitat at the location of sampling but also at nearby locations.
2. To address this, we extend methods from the Stochastic Partial Differential Equation (SPDE) method that is widely used in INLA, which approximates spatial correlations based on local diffusion over a finite-element mesh (FEM). We specifically introduce the sparse inverse-diffusion operator on a FEM, and apply this operator to covariates to efficiently calculate a spatially weighted average of local habitat that is then passed through pointwise basis-expansion to predict species densities. We show that this operator has several useful properties, i.e., conservation of mass, linear computational time with spatial resolution, and invariance to linear (scale and offset) transformations of covariates.
3. We test this covariate-diffusion method using a simulation experiment, and show that it can correctly recover a non-local environmental response while collapsing to a local (pointwise) response when warranted. We apply it to monitoring data for 25 bottom-associated fishes in the eastern Bering Sea and 20 bird species in the western United States. This application confirms that non-local responses in the eastern Bering Sea case study are parsimonious for 26 species-maturity combinations, while 18 collapse to the pointwise method. Estimates suggest that some species-maturity combinations avoid proximity to the continental slope, beyond what is predicted by local bathymetry in isolation. By contrast, in four of the 20 bird species is the diffused human population density covariate more parsimonious than the original covariate.
4. The covariate-diffusion method introduced here constitutes a fast and efficient approach to modelling non-local covariate effects. This flexible method may be useful in cases when covariates influence nearby population densities, for instance due to movement of the sampled species or its important biological or physical drivers.

## Introduction

Characterizing spatial patterns in the abundance of organisms in relation to environmental factors, and how that affects dynamics of ecological communities, is central to spatial ecology. At local scales, the abundance of species and demographic processes are shaped by both local habitat conditions, such as physical structure, competition and predation, and larger-scale processes (Menge and Olson 1990). The latter could refer to e.g., temperature and climate indices such as the North Atlantic Oscillation (Millon *et al.*2014), and to variables related to the dispersal pathways (Gómez-Pompa *et al.*1972, Jonsson *et al.*2016). Understanding how processes across spatial and temporal scales interact to shape species distribution and community structure is an important area of research in these times of rapid shifts in species distributions (Pinsky *et al.*2013, Roberts *et al.*2019, McCabe and Cobb 2021).

Species distribution models (SDMs) fitted to local occurrence, count, or biomass data are key tools in spatial ecology (Elith and Leathwick 2009). They can be used to quantify species’ distribution, abundance, and realized environmental niche and thereby be used to forecast range shifts (Liu *et al.*2023, Pinsky *et al.*2018). Over time, there has been a trend towards larger data sets over broader spatial and temporal scales (Rollinson *et al.*2021). This has led to increased power to detect effects and estimate functional relationships between covariates and responses, but also challenges related to non-stationarity. Non-stationarity here refers to the situation where the relationship between covariates and responses varies across space and/or time (Banerjee *et al.*2014, Rollinson *et al.*2021). In regression-based SDMs, which is the focus of this study, this form of nonstationarity can be accounted for by specifying effects of covariates that are allowed to evolve through time or vary in space (Hastie and Tibshirani 1993, Bartolino *et al.*2011, Thorson *et al.* 2023, Anderson *et al.*2024). Some examples include allowing the association of bottom-dwelling fishes with depth to change over time as they shift their distribution due to warming (English *et al.*2022), and allowing regional ocean condition indices to cause a density response that varies spatially (Lehodey *et al.*1997, Thorson 2019).

Another challenge related to spatial non-stationarity that has received less attention is the scale-dependence of covariates. Typically, local covariates are used in regression-based SDMs to infer the relationship between habitat covariates and the response variable. However, the true habitat an individual uses corresponds to the area it integrates via individual movement. Hence, for sessile species, local covariates may be warranted, but as species mobility increases, local-scale covariates would increasingly underestimate the habitat use in a typical scenario with a limited sample size. One could average covariates (and/or the response) prior to fitting the model to address that the relevant spatial scale that links covariates to the response is larger than the observation scale (e.g., Lindmark *et al.*2023, McKeon *et al.*2024), or evaluate multiple scales and find which leads to the best performance metrics (Bartolino *et al.*2012, Núñez-Riboni *et al.*2021). However, a limitation of this approach is that it is impossible to know the optimal scale of aggregation beforehand, and the scale resulting in the strongest effect does not necessarily mean it is the most relevant scale.

In this study, we introduce an approach that involves applying a diffusion operator to a covariate within the SPDE framework. This allows us to estimate the optimal spatial scale for computing a weighted average of a covariate, and can be thought of as a way to measure the effective “habitat area” that individuals are integrating via movement. Using simulation testing, we show how this covariate-diffusion model can correctly recover diffused covariate effects, or collapse to the raw covariate, when no covariate diffusion is present. We then apply the covariate-diffusion model to two real-world datasets on bottom-associated fishes and birds. We find that it is a parsimonious model for more than half the species-maturity combinations in the fish case study and four of the 20 bird species.

## Methods

### Covariate-diffusion

The Stochastic Partial Differential Equation (SPDE) method (Lindgren *et al.*2011) is widely used to define spatially correlated variables in statistical models in two continuous spatial dimensions. We briefly summarize the method here, before discussing how our covariate-diffusion model arises as a novel reuse of the underlying math.

At the highest level, the SPDE method seeks to specify a Gaussian random field (GRF) Z, where the value of this random field *z_s_* at a set of locations *s* ∈ *D* within a spatial domain *D* follows a Matérn covariance function

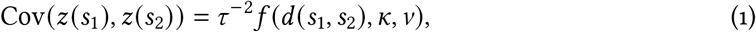

where *d* (*s*_1_, *s*_2_) is the distance between two locations, *f* (*d* (*s*_1_, *s*_2_), *κ, ν*) is the Matérn correlation function, *κ* is the decorrelation rate, *ν* is the smoothness parameter, and *τ* ^−2^ is the pointwise variance. This covariance function then allows the GRF to be evaluated at a fixed set of locations as a multivariate normal distribution

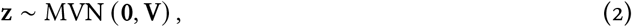

where V is the matrix of covariance among those locations. We could then calculate the value of the GRF *z*^*^ at a new location using bilinear interpolation, represented by a matrix A, *z*^*^ = Az.

In two-dimensional coordinates, and assuming that Matérn smoothness *ν* = 1, the SPDE method then approximates this GRF as a Gaussian Markov random field (GMRF) by specifying a sparse inverse-covariance (a.k.a. “precision” matrix) V^−1^ = Q

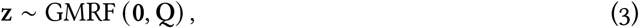

where evaluating the multivariate normal density function involves the precision matrix, and hence can be directly calculated from Q without matrix inversion. Importantly, the sparse precision matrix can also be constructed directly using the SPDE method

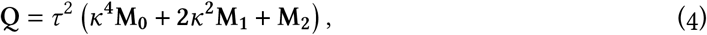

where M_0_ is a diagonal matrix, M_1_ has first-order adjacency within a triangulated mesh, and M_2_ has second-order adjacency. These three matrices are typically constructed by lower-level software (e.g., the R package fmesher Lindgren 2023), and fitting this model does not require advanced understanding of the model derivation. However, we here summarize the underlying theory to introduce our extension.

In particular, the SPDE approximation to a GMRF is derived by discretizing a diffusive process (the partial differential equation from the method’s name) and a stochastic “shock” *ϵ* as a simultaneous equation involving the realization z of our GMRF:

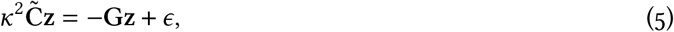

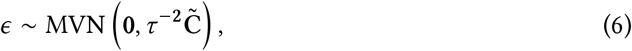

where 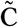 is a diagonal matrix and 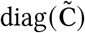 is the volume of the linear basis functions centered at each location and G is a sparse matrix representing the spatial overlap between basis functions (i.e., is zero for nonadjacent locations), as well as assumed boundary conditions for the SPDE process.

Adding Gz to both sides yields

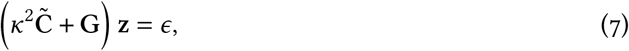

and then dividing the left-hand-side across and expressing as a GMRF yields

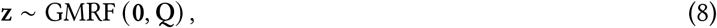

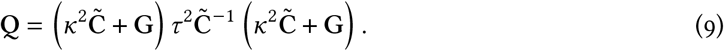

Multiplying out the quadratic form for the precision matrix then results in the original expression (Eq. 4), where 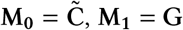, and 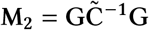.

Having re-iterated the diffusion process that underlies the SPDE precision matrix, we now define a diffusion matrix D (Supporting Information S1):

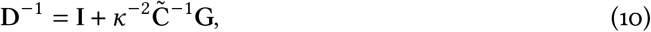

where the inverse-diffusion D^−1^ has the same sparsity as G, which follows first-order adjacency. This operator satisfies four desiderata:

1. *Conservation of mass*: Given a field approximated as vector z at the vertices of the finiteelement mesh, *N* evenly spaced locations s that cover the domain, and bilinear interpolation matrix A that projects to those *N* locations, we can approximate the average value of the field by predicting and then averaging across those locations 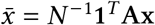. Pre-multiplying by the diffusion operator has (almost) no effect on this average mass, 1*^T^*Ax = 1*^T^*ADx, i.e., where the diffusion operator conserves the total value of z;
2. *Invariance to centering or scaling*: Given that we approximate diffusion using a linear operator, we can apply a linear transformation to any vector z^*^ = *a* + *b*z, and this will result in the same linear transformation of the diffused version Dz^*^ = *a* + *b*Dz. For example, if we measure temperature in Celcius at a set of sites, convert to Farhenheit, and then apply the diffusion operator, this will be equivalent to applying the diffusion operator and then converting to Farenheit;
3. *Invariance to geographic units*: If we multiply the geographic units by a constant (e.g., convert from kilometers to meters) with associated change in SPDE matrices (M_0_, M_1_, M_2_), and also divide the covariate diffusion rate *κ* by the same constant, then the diffusion operator remains unchanged;
4. *Efficient computation*: The diffusion matrix D is “dense” (i.e., values become small but remain nonzero even as distances become large), and hence the time to compute Dv scales as *S*^2^ where *S* is the number of sites (Supporting Information S2, Fig. S1). However, we can instead calculate Dv efficiently by first calculating a sparse LU decomposition of D^−1^ and then applying this to v. Doing so works directly with the sparse matrix D^−1^, and avoids computing or storing the dense matrix D.

We also note that this inverse-diffusion matrix operates across the entire domain of the FEM (which typically extends beyond the range of the data), and implicitly uses the Neumann (“reflective”) boundary condition (which is conventional when applying the SPDE approach). Covariate values must be specified for all FEM vertices (including boundary vertices), and future research could explore alternative options, for example, defining a Dirichlet (“absorptive”) boundary condition for covariate-diffusion at the precise edge of available data. However, we do not explore boundary conditions further here.

In the following, we therefore define a vector of covariate values x at the vertices of the finiteelement mesh, where the covariate x^*^ is interpolated at a new locations using the same interpolation matrix x^*^ = Ax. We then replace the covariate value *x*^*^ for sample *i* at location *s_i_* with its diffused value ADx, and use ADx to predict local densities in a species distribution model. The effect of applying the diffusion operator and its effect on the total mass of the covariate is visualized in Fig. 1. We then estimate parameter *κ* (used to construct diffusion matrix D) simultaneously with other regression coefficients representing habitat associations. As *κ* → ∞ in Eq. 10 then diffusion D^−1^ → I and the diffused covariate collapses on its local value Dx = x. Alternatively, as *κ* → 0 then Dx = *c*1 and the diffused covariate collapses on an constant value *c*. We are therefore interested in intermediate values of *κ* where Dx represents the impact of covariates within the neighborhood of a given location.

**Figure 1.**
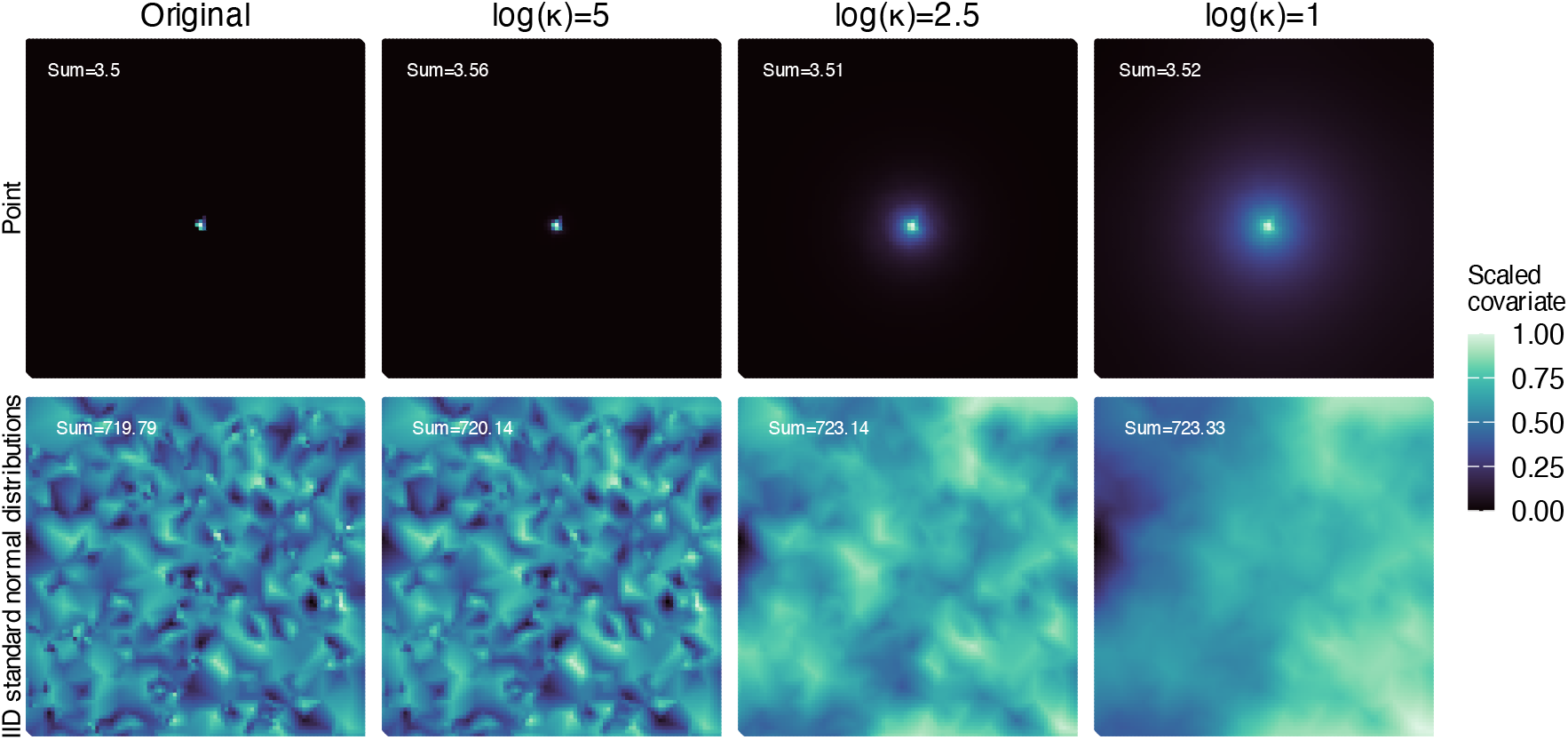
Applying the diffusion operator D when interpolating covariate x with interpolation matrix A to vertices of the finite-element mesh largely conserves the mass across different values of *κ* (i.e., ∑ Ax ≈ ∑ ADx). In the top row, the diffusion is visualized for a single central point and in the bottom row diffusion is applied to a vector of draws from IID standard normal distributions to visualize diffusion on the full covariate field. Columns correspond to different *κ* values, from large *κ* (low diffusion) to small *κ* (high diffusion). Note that the covariate values are scaled within each kappa scenario to visualize the diffusion; the number in the top left corresponds to the total mass of the covariate.

### Simulation testing

#### Testing the ability to recover diffusion with a simulation experiment

We developed a simulation experiment to explore the following questions: 1) how well the estimate for a diffused covariate could be recovered under varying observation error, 2) how often marginal AIC favoured the correct estimation model (diffusion or null model) in a self-and-cross experiment, 3) how well the diffusion parameters could be recovered under varying strengths of diffusion, and 4) how well the diffusion model can collapse to the null model (i.e., how well the diffusion model can match the null model estimates when data are simulated without diffusion). We simulated 200 datasets from a Poisson model with an observation-level random intercept in link space to allow for additional dispersion beyond the 1:1 mean-variance of the Poisson — a lognormal Poisson. Parameters were largely taken from a model fitted to counts of juvenile Pacific cod (*Gadus macrocephalus*) in a subsequent case study, with a scaled depth covariate (subtracting the mean and dividing by the standard deviation). Each data set contained 15592 spatially correlated observations, and for every dataset, a new GMRF was simulated. For a more detailed description of the models we refer to the northeastern Bearing Sea case study, see *Case studies*.

In the first exercise (questions 1 and 2), we generated data by simulating from models without and with covariate-diffusion. In the former, we set the intercept *β*_0_ to -0.4, the linear effect of the raw or diffused covariate *β _j_* to -2.4, the scalar of the precision matrix log(*τ*) to -1.4, and the decorrelation rate log(*κ_ω_*) to -0.8. For the diffusion model, we in addition set the strength of the diffusion log(*κ_X_*) to 2.5 (which corresponds to moderate diffusion). Henceforth, we refer to the diffusion parameter *κ_X_* (not to be confused with the decorrelation rate *κ_ω_*) as simply *κ*. In both operating models, we set the observation level standard deviation to values of 0.1, 1, and 2, and tested how well both models could return the true depth coefficient, and how often marginal AIC favoured the correct operating model.

In the second exercise (questions 3 and 4), we simulated data from a diffusion model to evaluate how well the true value of the diffused covariate could be retrieved, given varying strengths of the diffusion, and how often marginal AIC favoured the correct operating model. We used the same parameters for the diffusion model as above (*σ_η_* = 1), and set log(*κ*) to 1 (strong diffusion), 2.5 (moderate diffusion), and 5 (low diffusion). These levels of diffusion are illustrated in Fig. 1.

#### Speed comparisons

We also used simulation testing to compare fitting speeds between the two estimation models (null model and diffusion model). To this end, we simulated 1000, 10000, and 100000 spatially correlated observations following a Poisson distribution, with a spatial random effect, an intercept, and a predictor, using the R (R Core Team 2024) package sdmTMB (Anderson *et al.*2024). Next we compared the time to conduct numerical optimization and uncertainty quantification, across a range of mesh resolutions. To standardize and make timing benchmarks as comparable as possible across machines and operating systems, we used OpenBLAS (Xianyi *et al.*2012) and a single core.

#### Case studies

To illustrate a diffused covariate in practice, we also present two real-world case studies. In the first case study, we use count data for 20 species from the US Breeding Bird Survey (Sauer *et al.* 1997) in the western United States (westward of Wyoming, Colorado, Montana, and New Mexico) in 2019. We test if there is support for non-local effects of log human population density. This could, for instance, indicate a response to urbanization affecting the habitat quality. For example, outside densely populated areas there may still be large impacts of habitat due to infrastructure. We use marginal AIC to determine if the more complex covariate-diffusion model is supported. As a sensitivity test, we also calculated conditional AIC, following Zheng *et al.*(2024). The second case study is based on bottom trawl survey data from the northeastern Bering Sea in 2019, collected by the NOAA Alaska Fisheries Science Center using a fixed station design. Each trawled site contains information on catch in numbers of 44 combinations of species and maturation status. Here we test if there is support for non-local effects of sea floor depth. A diffused depth effect could, for instance, indicate a response to being near (but not actually on) the continental slope.

In both case studies, we modelled the counts at each site using a lognormal Poisson observation model and a log link

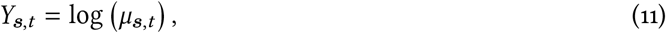

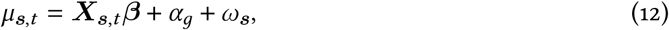

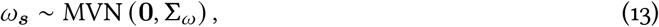

where *μ***_*s,t*_** represents the mean count, ***X*_*s,t*_** is the design matrix, *α_g_* is an observation-level random intercept, and *ω****_s_***represents spatial random effects drawn from a Gaussian Markov random field with inverse precision (i.e., covariance) matrix **Σ***_ω_* constrained by a Matérn covariance function.

We constructed finite element meshes using the function `fm_mesh_2d()`in the R package fmesher (Lindgren 2023), using a cutoff distance (minimum triangle edge length) of 0.1 degrees in the northeastern Bering Sea case study (1295 knots) and 1 degree in the Breeding Bird Survey (170 knots) (Supporting Information S2, Figs. S4–S3). Mesh construction is a complex topic, can impact parameter estimation, and involves a tradeoff between accuracy and estimation speed (Righetto *et al.*2020, Røste 2020, Commander *et al.*2022). The impact of mesh resolution on the diffusion model presented here is a topic of future research, but we expect mesh resolution will impact both the estimation of the diffusion process and the spatial random fields. Just as systems with less smooth spatial correlation will benefit from higher resolution meshes (Røste 2020), we expect that systems with more concentrated diffusion processes will require higher resolution meshes to accurately estimate the diffusion process. As a sensitivity test, we compared the outcomes of model selection across different mesh resolutions.

#### Estimation process

We fit the SPDE-based spatial models in the simulation experiment and the case studies using the R (R Core Team 2024) package TMB (Kristensen *et al.*2016) (version 1.9.17), with matrices in Eq. 5 constructed with the R package fmesher (Lindgren 2023). Parameter estimation is done via maximum marginal likelihood using the non-linear minimizer nlminb() in the R stats package (R Core Team 2024).

## Results

The covariate-diffusion estimation model is able to retrieve the true parameters accurately both when the underlying model (“operating model”) generating the data had covariate-diffusion and when it did not, since the diffusion model reverts to the sub model without covariate-diffusion as *κ* becomes large (Fig. 2). However when the operating model is a covariate-diffusion model, the null estimation model leads to biased parameter estimates (Fig. 2a). We also find that marginal AIC (mAIC) favours the covariate-diffusion model in *>*98% of iterations when the operating model is covariate-diffusion (Fig. 2a), and favours the null model in *>*89% of iterations when the operating model is null (Fig. 2b). Neither the ability to retrieve the true parameter estimate nor the assignment based on mAIC are affected by the observation error standard deviation (overdispersion) given the ranges tested here (Fig. 2). We also find that mAIC identifies the true operating model (diffusion) more frequently when the strength of the diffusion is medium or large (98–100% of iterations, respectively)(Fig. 3a–b). For low diffusion (Fig. 3c), mAIC favours the the null model in 90% of iterations. The covariate-diffusion model is able to retrieve the true parameter value on average regardless of the strength of the diffusion, but the spread of individual estimates is larger when the diffusion is stronger (Fig. 3a).

**Figure 2.**
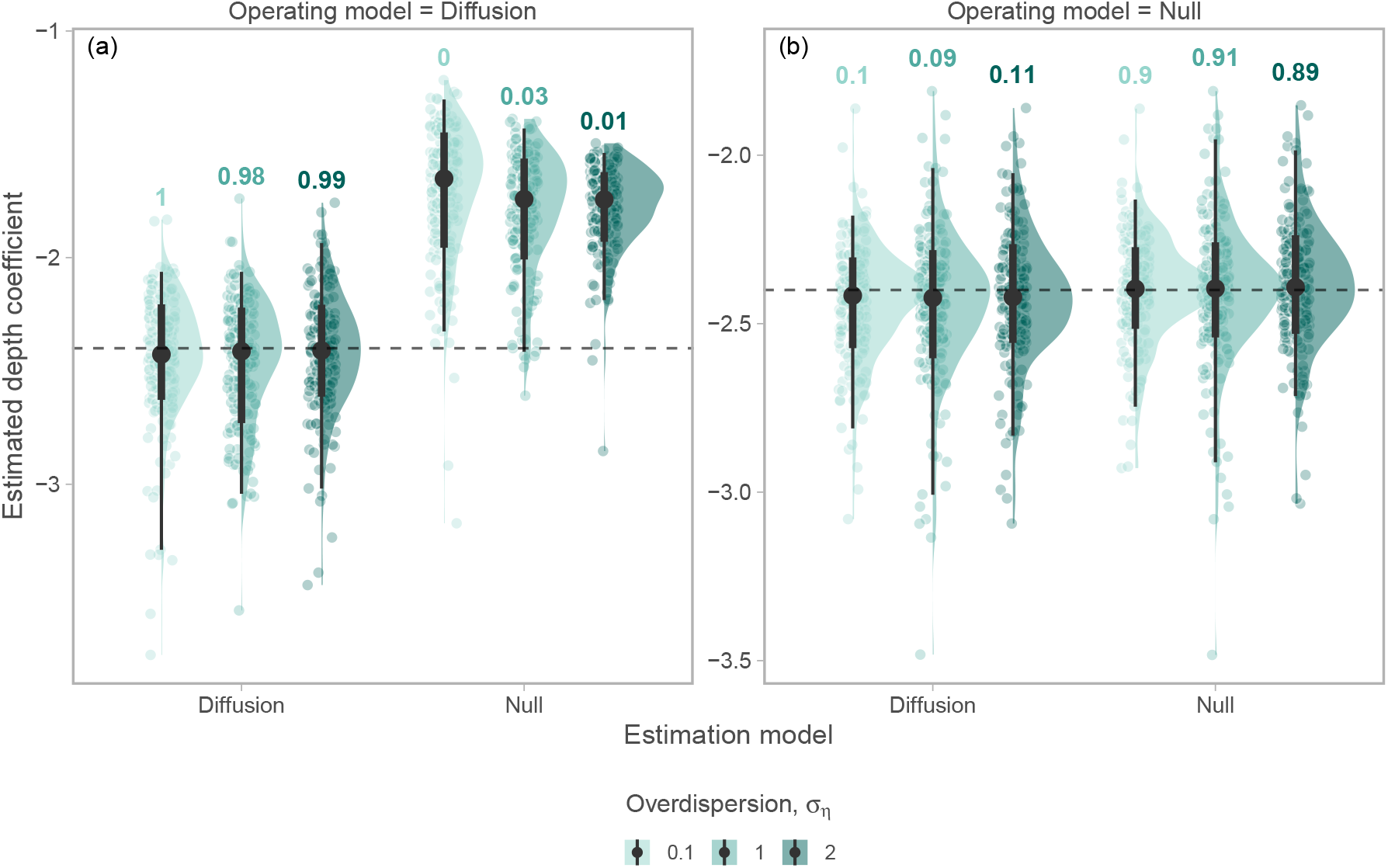
The diffusion model can recover diffused covariate effects and collapse to the null model in the absence of diffusion. Simulation testing the ability to recover the true estimated depth coefficient for diffusion and null operating models (left and right, respectively), for diffusion and null models (x-axis), for three levels of overdispersion, i.e., the rate at which the observation variance scales with the mean in excess of the Poisson *σ_η_* (color). The strength of the diffusion, log (*κ*), is set to 2.5 when the operating model is a diffusion model (left). Each point represents a fit from a simulated data set, black points and vertical lines correspond to the median, 50%, and 95% quantile range. Horizontal lines correspond to the true value. Numbers above vertical bars correspond to the proportion of simulated datasets (n=200) assigned to the estimation model (per value of *σ_η_*) based on marginal AIC.

**Figure 3.**
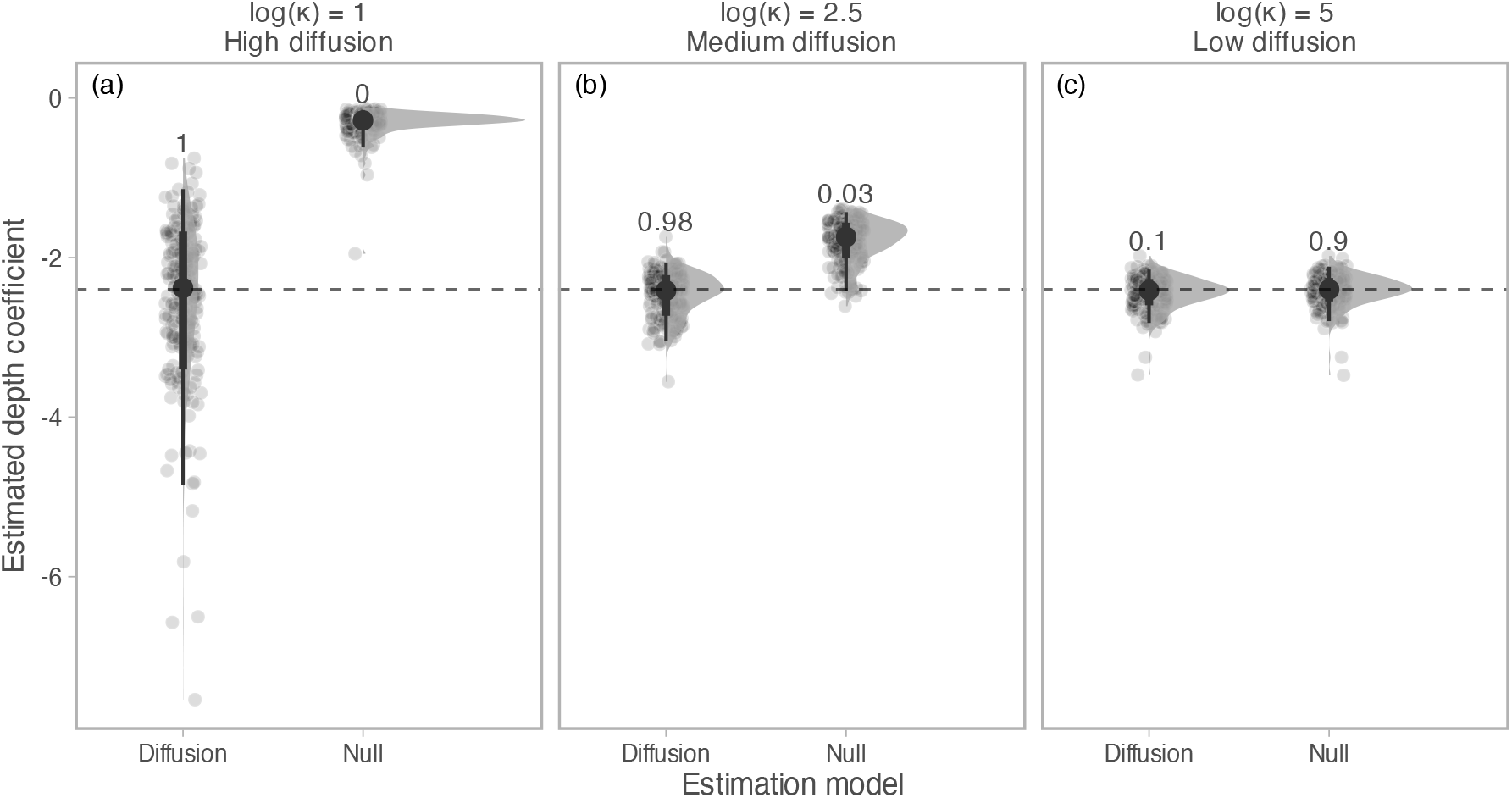
As the diffusion declines (*κ* increases, from left to right column), the difference between estimated depth coefficients from the diffusion and null models decreases (from left to right). Each point represents a fit from a simulated data set, black points and vertical lines correspond to the median, 50%, and 95% quantile range. Horizontal lines correspond to the true value. Numbers above vertical bars correspond to the proportion of simulated datasets (n=200) assigned to estimation model based on AIC.

Estimation time tends to increase with a diffusion model. The proportional increase in estimation time with a diffusion model increases with mesh resolution, but the rate of increase along the mesh axis depends on the number of observations. For larger data sets, the increase in estimation time with increased mesh resolution is less steep than with small data sets (Supporting Information S2, Fig. S2). Time to calculate standard errors is similar across estimation models (Supporting Information S2, Fig. S2).

Our case studies show that covariate diffusion is supported to varying degrees in both bird species and fish groups. In four of 20 bird species, covariate-diffusion is supported for the human population density covariate as they have ΔmAIC *>* 2 (Fig. 4a). In contrast, we find support for covariate diffusion of a quadratic depth effect in 26 out of 44 species-maturity combinations in the eastern Bering Sea case study on fishes (Fig. 4b). In both case studies, a ΔmAIC = −2 indicates that the covariate-diffusion and the null model have the same marginal log likelihood, and the correlation between the raw and diffused covariate approaches 1 (Fig. 4). When the same comparison is done with conditional AIC (cAIC) instead, which generally penalizes complexity more heavily than marginal AIC since it includes a correction for the number of random effects estimated, 18 of the fish species and 0 bird species show support for covariate diffusion in our examples (Supporting Information S2, Fig. S5). We also find that these results are relatively consistent across mesh-resolutions in the Eastern Bering sea case study. For example, in 64% of species, mAIC favors the same model across 3 different mesh resolutions (Supporting Information S2, Fig. S6).

**Figure 4.**
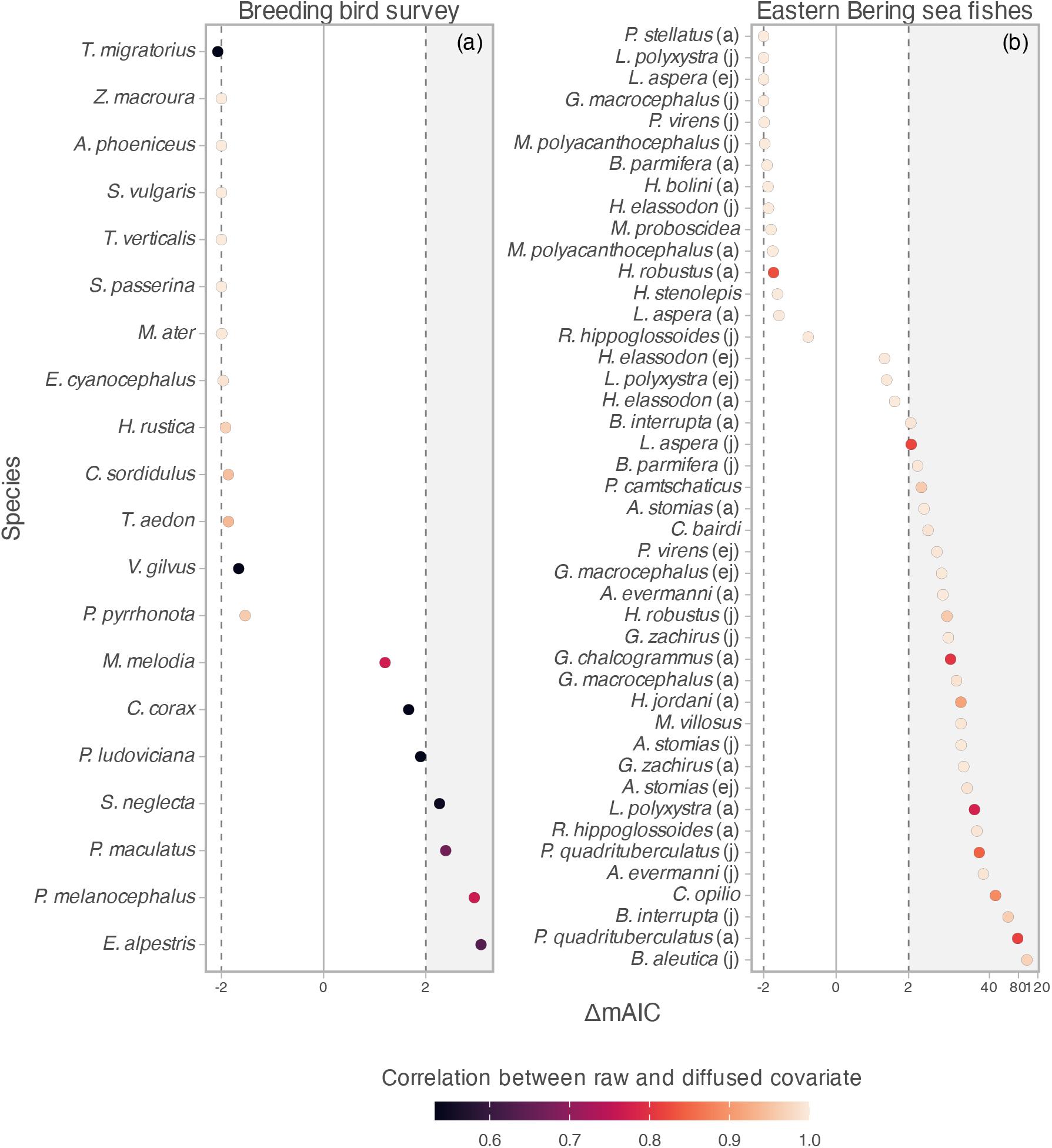
Marginal AIC favours covariate-diffusion in more than half of fishes and four bird species. The points depict delta marginal AIC between the null and diffusion model, where positive values indicate support for the diffusion model and negative values indicate support for the null model. Point colours correspond to the correlation between the raw and the diffused covariate. Points in the grey rectangle have ΔmAIC >2, indicating strong support for the diffusion model. Points within the two vertical dashed lines have inconclusive ΔmAIC results. Letters in brackets in the Eastern Bering sea fish case study refers to the life stage (j=juvenile, a=adult, ej = early juvenile). Note the x-axis is fourth-root power transformed.

A lower correlation between the diffused and raw covariate is typically found in species where the covariate-diffusion model is supported (Fig. 4). For example, in the Breeding Bird case, the covariate-diffusion model is not supported for the common starling (*Sturnus vulgaris*; top row) and the diffused covariate (middle column) is nearly identical to the original covariate (left column), while for black-headed grosbeak (*Pheucticus melanocephalus*; bottom row) the human population density covariate is smoothed with a strong diffusion (log(*κ*) = −1.09) (Fig. 5b). Similarly for the example fishes from the northeastern Bearing Sea case, in adult starry flounder (*Platichthys stellatus*), the depth covariate from the covariate-diffusion model is nearly identical to the raw covariate, while for capelin (*Mallotus villosus*), the diffusion is moderate to strong (log *κ* = 2.76), and the covariate exhibits a smoother pattern.

**Figure 5.**
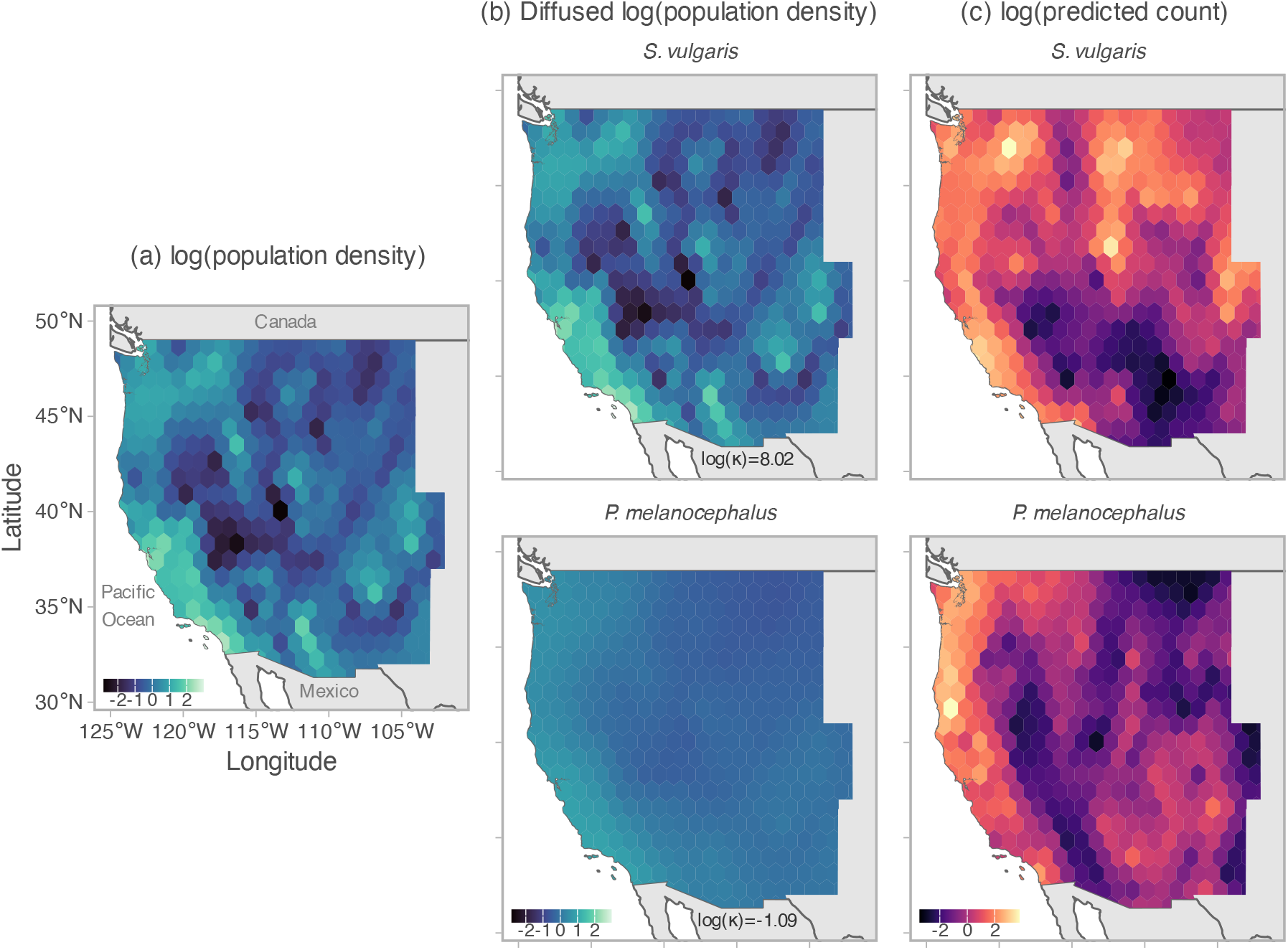
Human population density covariate, a diffused version of the covariate, and predicted counts from the breeding bird case study. Panel (a) depicts the raw human population density covariate. Panel (b) depicts the diffused covariate for two species with contrasting support for diffusion. Common starling (*Sturnus vulgaris*) in the top row does not show support for the diffused covariate and the diffusion is estimated to be small whereas black-headed grosbeak (*Pheucticus melanocephalus*) in the bottom row shows strong support for the diffused covariate and has a relatively strong estimated diffusion. The strength of the diffusion (log (*κ*)) is shown towards the bottom of the (b) panels, where a low value indicates strong diffusion. Panel (c) depicts the predicted log counts for the two species.

**Figure 6.**
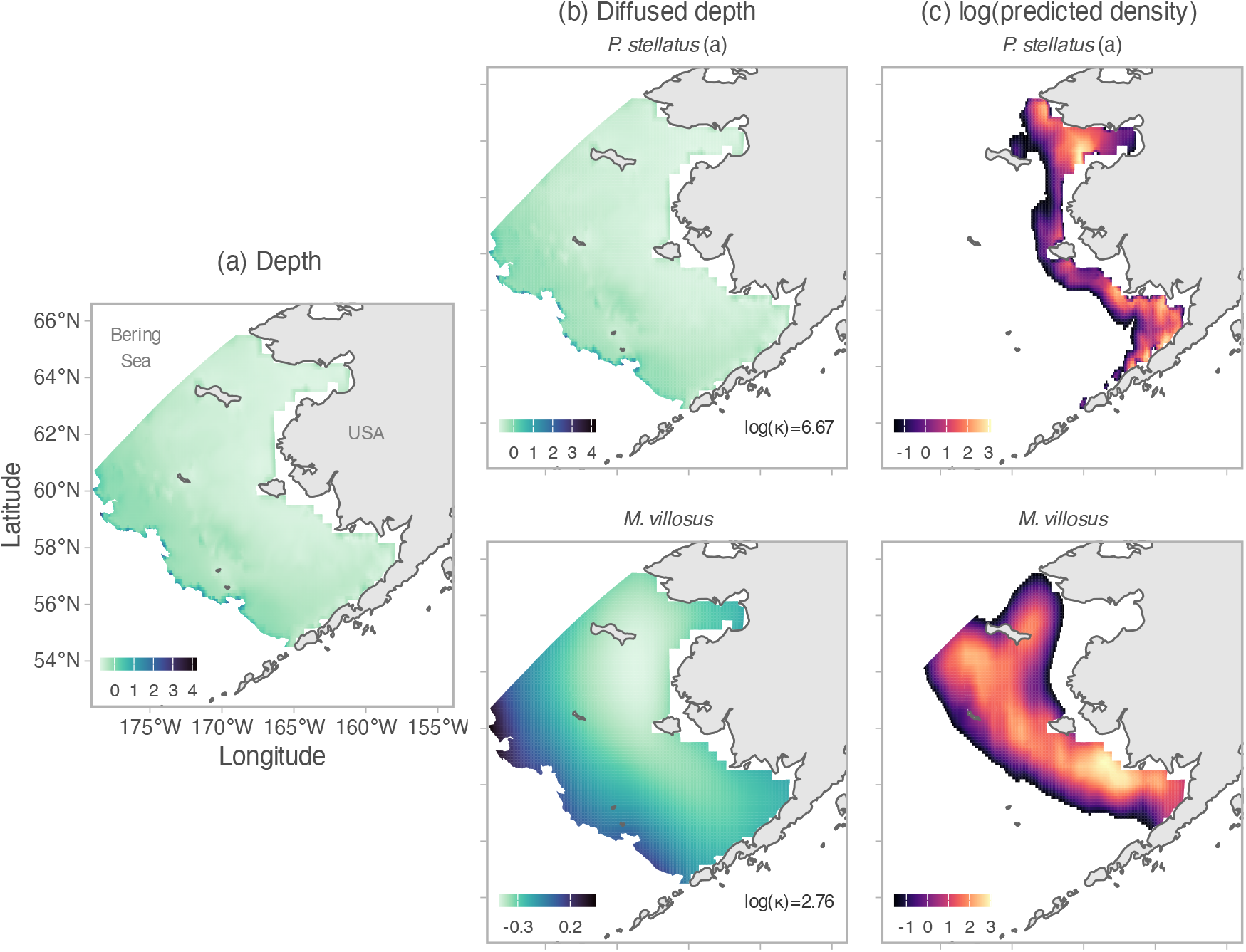
Bottom depth covariate, a diffused version of the covariate, and predicted counts from the northeastern Bering sea bottom trawl data. Panel (b) depicts the diffused covariate for two species with contrasting support for diffusion. Adult starry flounder (*Platichthys stellatus*) in the top row does not support the diffusion model and the diffused covariate is similar to the raw covariate, while capelin (*Mallotus villosus*) in the bottom row shows strong support for the diffusion model. The strength of the diffusion (log (*κ*)) is shown in the bottom-right corner of the (b) panels; a low value indicates strong diffusion. Panel (c) depicts the predicted log counts for the two species (values *<* 1% of the maximum density are omitted for visualization).

For several species-maturity combinations in the fish case study, there is a notable difference in the partial effect of depth on density (Fig. S7). However, the covariate-diffusion model and the null model often generate similar predictions, even in cases of strong diffusion, presumably because the spatial random effects can change between the models (Supporting Information S2, Figs. S8, S9).

## Discussion

We have introduced a sparse inverse-diffusion operator based on the SPDE method, which can be used to efficiently model non-local covariate effects such as to approximate the effective habitat area that individuals integrate via movement. Specifically, when applied to a covariate, this operator calculates a spatially weighted average covariate given the estimated range of the diffusion processes. With simulation testing, we have demonstrated that the diffusion model can correctly identify the underlying processes model and estimate the density response to the diffused covariate.

We then tested the approach on spatial models fitted to datasets for bird and fishes. Covariatediffusion was more parsimonious than the null model for only two of 20 bird species, but for a majority of species-maturity combinations in fishes. As an example of interpretation, for some species-maturity combinations in the fish case study, the partial effect of depth was smaller for the covariate diffusion model than the null model near the continental shelf slope suggesting that these groups avoid these habitats despite being of similar depths to other more inshore areas. Hence, our approach could aid generating hypothesis as to what drives non-stationary in across space, which is important for improving large-scale species distribution modeling (Rollinson *et al.*2021).

Covariate-diffusion could result from any ecological teleconnections such that local ecological properties are influenced by patterns happening at a broader scale. For example, fish move over time and therefore their body condition (how plump they are given their length) may be affected by the combination of habitat and spatially varying prey they encounter over their lifetime (Lindmark *et al.*2023). Covariate diffusion could represent how this broader scale of conditions the fish moved through might affect their body condition. Alternatively, a species may be stationary with environmental process changing around them. For example, the number of eggs produced by sessile clams may be influenced by environmental conditions as ocean currents move water past the clams. Covariate diffusion could represent how this broader scale of experienced environment might affect clam fecundity. Future research could extend our approach by estimating diffusion that differs based on direction (i.e., extracting the diffusion kernel from the SPDE method given geometric anisotropy), or by incorporating covariate effects that decay across space and persist into future times (i.e., extracting the diffusion kernel from a diffusion-enhanced GMRF, see Lindgren *et al.*(2023)). Furthermore, the inverse-diffusion operator should remain sparse (and therefore computational efficient) in these and other cases.

We observe that predicted densities from the diffusion model and the null model tend to be similar. While both the covariate and the estimate of its coefficient change when the diffusion model is applied and supported, the predicted counts do not substantially differ between the two models, partly because the spatial random effects also change. Which model to use then depends on the objectives of the analysis—whether it is to learn about ecologically relevant scales of covariates and non-local effects or if a model that generates similar predictions by placing additional variation in the spatial random effects will suffice. Since we have also shown that the diffusion model can revert to a non-diffused model in the absence of diffusion, the diffusion model can be applied at little cost even when it is not known a priori whether diffusion is supported.

We recommend three topics for future research. The first is to augment our covariate-diffusion model by incorporating advection, i.e., where local densities respond to environmental conditions that are centered on a location that is geographically distant. This “covariate-advection” is feasible using the SPDE method (Clarotto *et al.*2023) and would presumably represent advective movement, e.g., where densities during summer sampling respond to habitat conditions in a winter habitat. Secondly, we note that covariate-diffusion collapses to an index of regionally averaged conditions as diffusion becomes large. In this case, fitting a spatially varying coefficient (SVC) (Hastie and Tibshirani 1993, Gelfand *et al.*2003) response to the diffused covariate across multiple years would allow a wide range of model behaviors, from a stationary and local response to a non-stationary response to a regional climate index. Lastly, future research could investigate best practices for model selection and mesh construction comparing covariate-diffusion models, as our results suggest some sensitivity to choice of selection criteria and mesh resolution. It is also important that these practices are tied together with objectives — whether prioritizing predictive accuracy or ecological inference.

We note several drawbacks to the covariate-diffusion approach. First, the approach replaces the high-resolution covariate measured at each unique location with an interpolated value that is defined at each vertex of the finite-element mesh. This mesh can be defined at a high resolution, but still requires some loss of fine-scale variation. Second, although computationally efficient due to working with the sparse inverse-diffusion matrix, the approach is still more computationally intensive than fitting a model without covariate diffusion. Third, the model requires users to define covariate values not just for the location of samples, but at all locations across a given domain. This results in a more-complex user interface than the regression models typically used for SDMs and will therefore require some consideration before integrating into GMRF- and TMB-based SDM software such as sdmTMB (Anderson *et al.*2024) or tinyVAST (Thorson *et al.*2024).

Despite these drawbacks, we conclude that covariate-diffusion using the SPDE method is computationally efficient, statistically performant, and ecologically important for a wide range of species. We therefore recommend that ecologists estimate non-local habitat responses across the wide range of studies applying SDMs.

## Acknowledgements

We thank everyone involved in the collection, processing and collation of trawl survey and bird count data. We thank C. Freshwater and two anonymous reviewers for comments that improved this paper. The authors have no conflicts of interests to declare.

## References

Anderson, S.C., Ward, E.J., English, P.A., Barnett, L.A.K. and Thorson, J.T. (2024) sdmTMB: An R package for fast, flexible, and user-friendly generalized linear mixed effects models with spatial and spatiotemporal random fields. bioRxiv 2022.03.24.485545. URL 10.1101/2022.03.24.485545.

Banerjee, S., Carlin, B.P. and Gelfand, A.E. (2014) Hierarchical Modeling and Analysis for Spatial Data. 2 edition. Chapman and Hall/CRC, New York.

Bartolino, V., Ciannelli, L., Bacheler, N.M. and Chan, K.S. (2011) Ontogenetic and sex-specific differences in density-dependent habitat selection of a marine fish population. Ecology 92, 189–200. URL https://onlinelibrary.wiley.com/doi/abs/10.1890/09-1129.1. eprint: https://onlinelibrary.wiley.com/doi/pdf/10.1890/09-1129.1.

Bartolino, V., Ciannelli, L., Spencer, P., Wilderbuer, T.K. and Chan, K.S. (2012) Scale-dependent detection of the effects of harvesting a marine fish population. Marine Ecology Progress Series 444, 251–261. URL https://www.int-res.com/abstracts/meps/v444/p251-261/.

Clarotto, L., Allard, D., Romary, T. and Desassis, N. (2023) The SPDE approach for spatio-temporal datasets with advection and diffusion.

Commander, C.J.C., Barnett, L.A.K., Ward, E.J., Anderson, S.C. and Essington, T.E. (2022) The shadow model: How and why small choices in spatially explicit species distribution models affect predictions. PeerJ 10, e12783.

Elith, J. and Leathwick, J.R. (2009) Species Distribution Models: Ecological Explanation and Prediction Across Space and Time. Annual Review of Ecology, Evolution, and Systematics 40, 677–697. URL https://www.annualreviews.org/content/journals/10.1146/annurev.ecolsys.110308.120159. Publisher: Annual Reviews.

English, P.A., Ward, E.J., Rooper, C.N., Forrest, R.E. et al. (2022) Contrasting climate velocity impacts in warm and cool locations show that effects of marine warming are worse in already warmer temperate waters. Fish and Fisheries 23, 239–255. URL https://onlinelibrary.wiley.com/doi/abs/10.1111/faf.12613. eprint: https://onlinelibrary.wiley.com/doi/pdf/10.1111/faf.12613.

Gelfand, A.E., Kim, H.J., Sirmans, C.F. and Banerjee, S. (2003) Spatial modeling with spatially varying coefficient processes. Journal of the American Statistical Association 98, 387–396.

Gómez-Pompa, A., Vázquez-Yanes, C. and Guevara, S. (1972) The Tropical Rain Forest: A Nonrenewable Resource. Science 177, 762–765. URL https://www.science.org/doi/10.1126/science.177.4051.762.

Hastie, T. and Tibshirani, R. (1993) Varying-Coefficient Models. Journal of the Royal Statistical Society. Series B (Methodological) 55, 757–796.

Jonsson, P.R., Corell, H., André, C., Svedäng, H. and Moksnes, P.O. (2016) Recent decline in cod stocks in the North Sea–Skagerrak–Kattegat shifts the sources of larval supply. Fisheries Oceanography 25, 210–228. URL https://onlinelibrary.wiley.com/doi/abs/10.1111/fog.12146. eprint: https://onlinelibrary.wiley.com/doi/pdf/10.1111/fog.12146.

Kristensen, K., Nielsen, A., Berg, C.W., Skaug, H. and Bell, B.M. (2016) TMB: Automatic Differentiation and Laplace Approximation. Journal of Statistical Software 70, 1–21. URL https://www.jstatsoft.org/index.php/jss/article/view/v070i05. xNumber: 1.

Lehodey, P., Bertignac, M., Hampton, J., Lewis, A. and Picaut, J. (1997) El Niño Southern Oscillation and tuna in the western Pacific. Nature 389, 715–718. URL https://www.nature.com/articles/39575.

Lindgren, F. (2023) fmesher: Triangle Meshes and Related Geometry Tools. URL https://CRAN.R-project.org/package=fmesher. R package version 0.1.5.

Lindgren, F., Bakka, H., Bolin, D., Krainski, E. and Rue, H. (2023) A diffusion-based spatio-temporal extension of Gaussian Matérn fields. URL http://arxiv.org/abs/2006.04917. 2006.04917 [stat].

Lindgren, F., Rue, H. and Lindström, J. (2011) An explicit link between Gaussian fields and Gaussian Markov random fields: the stochastic partial differential equation approach. Journal of the Royal Statistical Society: Series B (Statistical Methodology) 73, 423–498. URL http://onlinelibrary.wiley.com/doi/10.1111/j.1467-9868.2011.00777.x/abstract.

Lindmark, M., Anderson, S.C., Gogina, M. and Casini, M. (2023) Evaluating drivers of spatiotemporal variability in individual condition of a bottom-associated marine fish, Atlantic cod (Gadus morhua). ICES Journal of Marine Science 80, 1539–1550. URL 10.1093/icesjms/fsad084.

Liu, O.R., Ward, E.J., Anderson, S.C., Andrews, K.S. et al. (2023) Species redistribution creates unequal outcomes for multispecies fisheries under projected climate change. Science Advances 9, eadg5468. URL https://www.science.org/doi/full/10.1126/sciadv.adg5468. Publisher: American Association for the Advancement of Science.

McCabe, L.M. and Cobb, N.S. (2021) From Bees to Flies: Global Shift in Pollinator Communities Along Elevation Gradients. Frontiers in Ecology and Evolution 8. URL https://www.frontiersin.org/journals/ecology-and-evolution/articles/10.3389/fevo.2020.626124/full. Publisher: Frontiers.

McKeon, C.M., Buckley, Y.M., Moriarty, M., Lundy, M. and Kelly, R. (2024) Increased signal of fishing pressure on community life-history traits at larger spatial scales. Global Ecology and Biogeography 33, e13815. URL https://onlinelibrary.wiley.com/doi/abs/10.1111/geb.13815. eprint: https://onlinelibrary.wiley.com/doi/pdf/10.1111/geb.13815.

Menge, B.A. and Olson, A.M. (1990) Role of scale and environmental factors in regulation of community structure. Trends in Ecology & Evolution 5, 52–57. URL https://www.cell.com/trends/ecology-evolution/abstract/0169-5347(90)90048-I. Publisher: Elsevier.

Millon, A., Petty, S.J., Little, B., Gimenez, O., Cornulier, T. and Lambin, X. (2014) Dampening prey cycle overrides the impact of climate change on predator population dynamics: a long-term demographic study on tawny owls. Global Change Biology 20, 1770– 1781. URL https://onlinelibrary.wiley.com/doi/abs/10.1111/gcb.12546. eprint: https://onlinelibrary.wiley.com/doi/pdf/10.1111/gcb.12546.

Núñez-Riboni, I., Akimova, A. and Sell, A.F. (2021) Effect of data spatial scale on the performance of fish habitat models. Fish and Fisheries 22, 955–973. URL https://onlinelibrary.wiley.com/doi/abs/10.1111/faf.12563. eprint: https://onlinelibrary.wiley.com/doi/pdf/10.1111/faf.12563.

Pinsky, M.L., Reygondeau, G., Caddell, R., Palacios-Abrantes, J., Spijkers, J. and Cheung, W.W.L. (2018) Preparing ocean governance for species on the move. Science 360, 1189–1191. URL https://www.science.org/doi/full/10.1126/science.aat2360. Publisher: American Association for the Advancement of Science.

Pinsky, M.L., Worm, B., Fogarty, M.J., Sarmiento, J.L. and Levin, S.A. (2013) Marine Taxa Track Local Climate Velocities. Science 341, 1239–1242. URL https://www.science.org/doi/full/10.1126/science.1239352. Publisher: American Association for the Advancement of Science.

R Core Team (2024) R: A Language and Environment for Statistical Computing. R Foundation for Statistical Computing, Vienna, Austria. URL https://www.R-project.org/.

Righetto, A.J., Faes, C., Vandendijck, Y. and Jr, P.J.R. (2020) On the choice of the mesh for the analysis of geostatistical data using R-INLA. Communications in Statistics - Theory and Methods 49, 203–220.

Roberts, C.P., Allen, C.R., Angeler, D.G. and Twidwell, D. (2019) Shifting avian spatial regimes in a changing climate. Nature Climate Change 9, 562–566. URL https://www.nature.com/articles/s41558-019-0517-6. Publisher: Nature Publishing Group.

Rollinson, C.R., Finley, A.O., Alexander, M.R., Banerjee, S. et al. (2021) Working across space and time: nonstationarity in ecological research and application. Frontiers in Ecology and the Environment 19, 66–72. URL https://esajournals.onlinelibrary.wiley.com/doi/10.1002/fee.2298.

Røste, J. (2020) The Importance of Mesh Resolution When Using the SPDE Approach. MSc Thesis, Norwegian University of Science and Technology.

Sauer, J.R., Hines, J.E., Gough, G., Thomas, I. and Peterjohn, B.G. (1997) The north american breeding bird survey results and analysis. Technical report, Eastern Ecological Science Center, Laurel, MD.

Thorson, J.T. (2019) Measuring the impact of oceanographic indices on species distribution shifts: The spatially varying effect of cold-pool extent in the eastern Bering Sea. Limnology and Oceanography 64, 2632–2645. URL https://aslopubs.onlinelibrary.wiley.com/doi/abs/10.1002/lno.11238. eprint: https://aslopubs.onlinelibrary.wiley.com/doi/pdf/10.1002/lno.11238.

Thorson, J.T., Anderson, S.C., Goddard, P. and Rooper, C.N. (2024) tinyVAST: R package with an expressive interface to specify lagged and simultaneous effects in multivariate spatio-temporal models.

Thorson, J.T., Barnes, C.L., Friedman, S.T., Morano, J.L. and Siple, M.C. (2023) Spatially varying coefficients can improve parsimony and descriptive power for species distribution models. Ecography 2023, e06510. URL https://onlinelibrary.wiley.com/doi/abs/10.1111/ecog.06510. eprint: https://onlinelibrary.wiley.com/doi/pdf/10.1111/ecog.06510.

Xianyi, Z., Qian, W. and Yunquan, Z. (2012) Model-driven level 3 BLAS performance optimization on Loongson 3A processor. In: 2012 IEEE 18th International Conference on Parallel and Distributed Systems. pp. 684–691.

Zheng, N., Cadigan, N. and Thorson, J.T. (2024) A note on numerical evaluation of conditional Akaike information for nonlinear mixed-effects models. URL http://arxiv.org/abs/2411.14185. 2411.14185 [stat].

