## Supporting Information S1 for "Estimating scale-dependent covariate responses using two-dimensional diffusion derived from the SPDE method"

In the main text (Eq. 5), we introduced the simultaneous-equation form for the SPDE method:

$$\kappa^2 \tilde{\mathbf{C}} \mathbf{z} = -\mathbf{G}\mathbf{z} + \epsilon, \tag{S1}$$

where $\tilde{\mathbf{C}}$ and $\mathbf{G}$ are calculated from a finite-element mesh over a spatial domain. Solving for $\mathbf{z}$ as a Gaussian Markov random field then results in the widely used expression for the sparse precision matrix (Eq. 4). We then claimed without derivation that this same simultaneous equation can result in the sparse inverse-diffusion operator (Eq. 10):

$$\mathbf{D}^{-1} = \mathbf{I} + \kappa^{-2} \tilde{\mathbf{C}}^{-1} \mathbf{G}, \tag{S2}$$

where $\mathbf{x}^* = \mathbf{D}\mathbf{x}$ then serves as a diffusive covariate that integrates locally across the original covariate $\mathbf{x}$. Here, we provide a brief sketch of this derivation.

We start by defining the ordinary differential equation representing diffusive movement with instantaneous diffusion rate $\mathbf{D}'$:

$$\frac{\partial}{\partial t}\mathbf{x} = \mathbf{D}'\mathbf{x}, \tag{S3}$$

where $\mathbf{x}^*$ is calculated by integrating this ordinary differential equation system. In this interpretation, the covariate $\mathbf{x}$ undergoes one "time step" of integrating the instantaneous diffusion rate $\mathbf{D}'$ to yield the spatially autocorrelated variable $\mathbf{x}^*$. The integral can be approximated using an implicit first-order Euler method with unit time step:

$$\mathbf{x}^* = \mathbf{D}'\mathbf{x}^* + \mathbf{x}. \tag{S4}$$

Rearranging Eq. S4 gives

$$(\mathbf{I} - \mathbf{D}')\mathbf{x}^* = \mathbf{x}, \tag{S5}$$

which can be solved for

$$\mathbf{x}^* = (\mathbf{I} - \mathbf{D}')^{-1}\mathbf{x} = \mathbf{D}\mathbf{x}. \tag{S6}$$

such that the implicit first-order Euler method approximates $\mathbf{D}$ as $(\mathbf{I} - \mathbf{D}')^{-1}$.

To make use of this, we have to reformulate Eq. S1 so that it resembles Eq. S4. To do so, we take Eq. S1 and multiply $\kappa^{-2}\tilde{\mathbf{C}}^{-1}$ across both sides of Eq. S1, which yields:

$$\mathbf{z} = -\kappa^{-2}\tilde{\mathbf{C}}^{-1}\mathbf{G}\mathbf{z} + \kappa^{-2}\tilde{\mathbf{C}}^{-1}\epsilon. \tag{S7}$$

We then define the following substitutions:

$$\mathbf{D}' = -\kappa^{-2}\tilde{\mathbf{C}}^{-1}\mathbf{G} \tag{S8}$$

$$\mathbf{x} = \kappa^{-2}\tilde{\mathbf{C}}^{-1}\epsilon \tag{S9}$$

$$\mathbf{x}^* = \mathbf{z}, \tag{S10}$$

which translate Eq. S7 to Eq. S4. In a sense, we use the original SPDE method, but replace a transformation of the white-noise $\epsilon$ with our covariate, and use the finite-element matrices to define the instantaneous diffusion rate.

From Eq. S8, we see that:

$$\mathbf{D} = (\mathbf{I} - \mathbf{D}')^{-1} = \left(\mathbf{I} + \kappa^{-2}\tilde{\mathbf{C}}^{-1}\mathbf{G}\right)^{-1}. \tag{S11}$$

Therefore, rather than forming the dense diffusion matrix $\mathbf{D}$, we can compute $\mathbf{x}^* = \mathbf{D}\mathbf{x}$ by solving the linear system:

$$\left(\mathbf{I} + \kappa^{-2}\tilde{\mathbf{C}}^{-1}\mathbf{G}\right)\mathbf{x}^* = \mathbf{x} \tag{S12}$$

using sparse matrix factorization methods such as the sparse-LU decomposition of the matrix $\mathbf{D}^{-1} = \mathbf{I} + \kappa^{-2}\tilde{\mathbf{C}}^{-1}\mathbf{G}$.
