## Supporting Information S2 for "Estimating scale-dependent covariate responses using two-dimensional diffusion derived from the SPDE method"

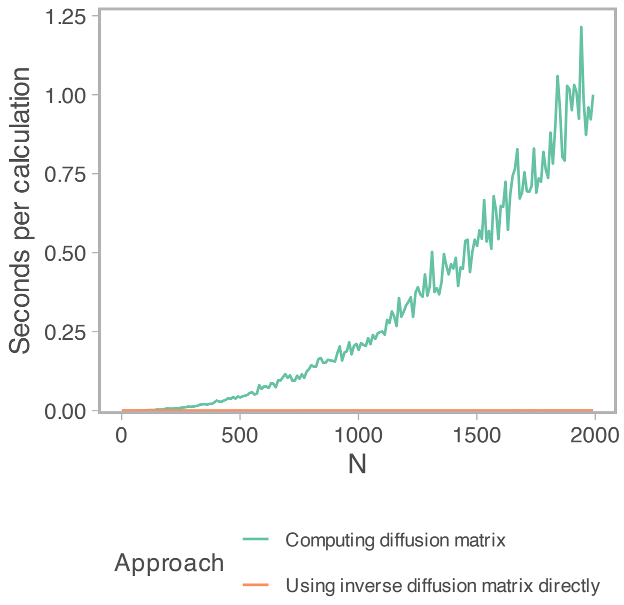


Figure S1: Comparison of time to compute $\mathbf{D}^{-1}\mathbf{x}$ using a sparse LU decomposition (orange) or by constructing the dense matrix $\mathbf{D}$ (green) given number of sites (x-axis).

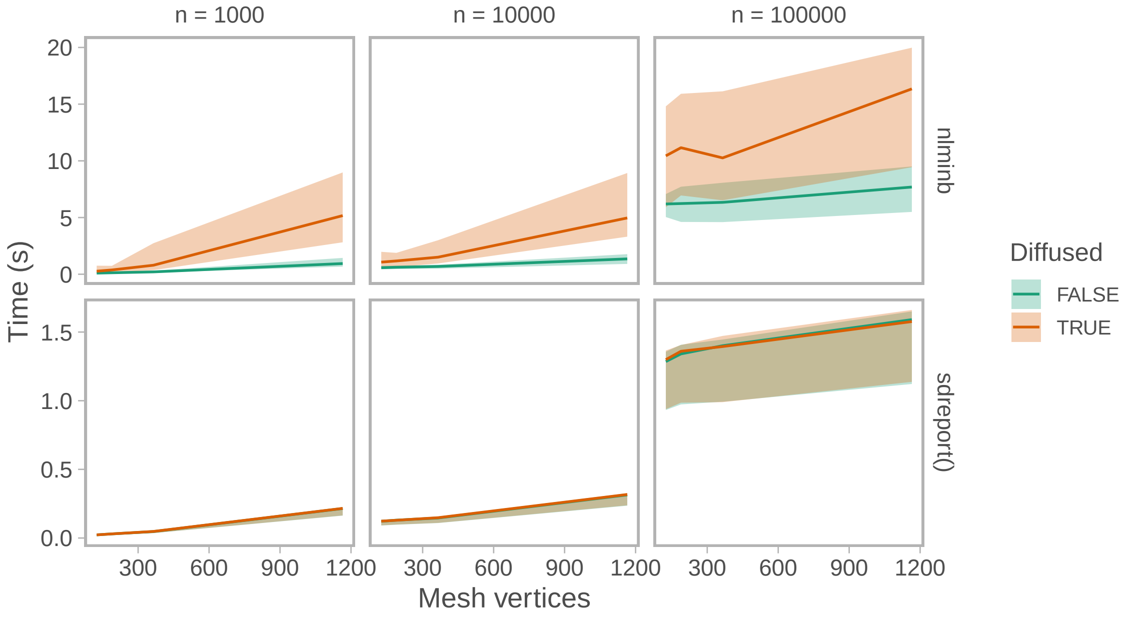


Figure S2: Comparison of time in seconds to estimate parameters (top row) and to calculate standard errors (bottom row) for a diffused model (orange), and a non-diffused model (green). Data were simulated from a Poisson distribution (left: n=1000, middle: n=10000, right: n=100000), and the log-linked Poisson GLMM estimated an intercept, the covariate diffusion rate and a slope, and a spatial spatial random field including two fixed effects defining its correlation function. The speed test was replicated across a range of SPDE mesh resolutions (x-axis). Lines depict the median and the ribbon encompasses the $5^{th}$–$95^{th}$ percentile across 50 random seeds.

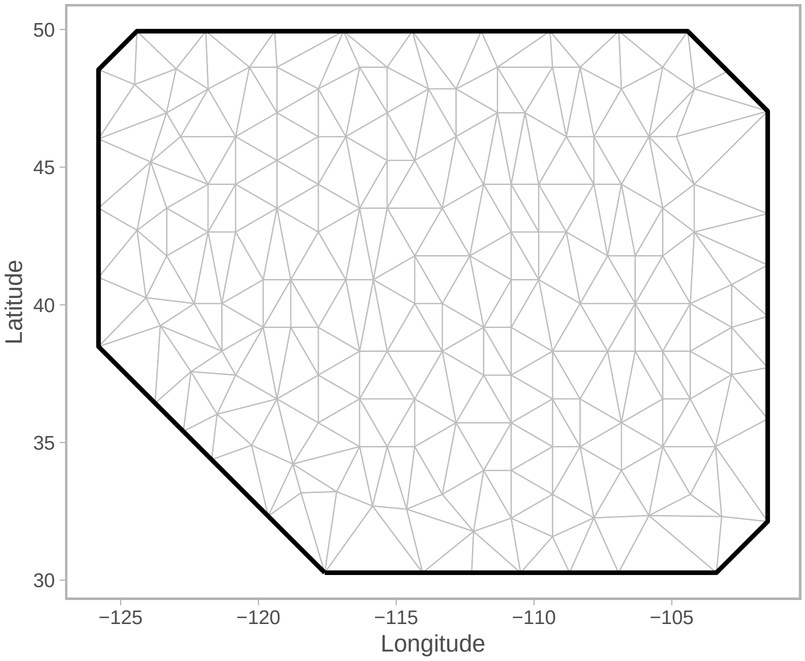


Figure S3: SPDE mesh for the breeding bird case study.

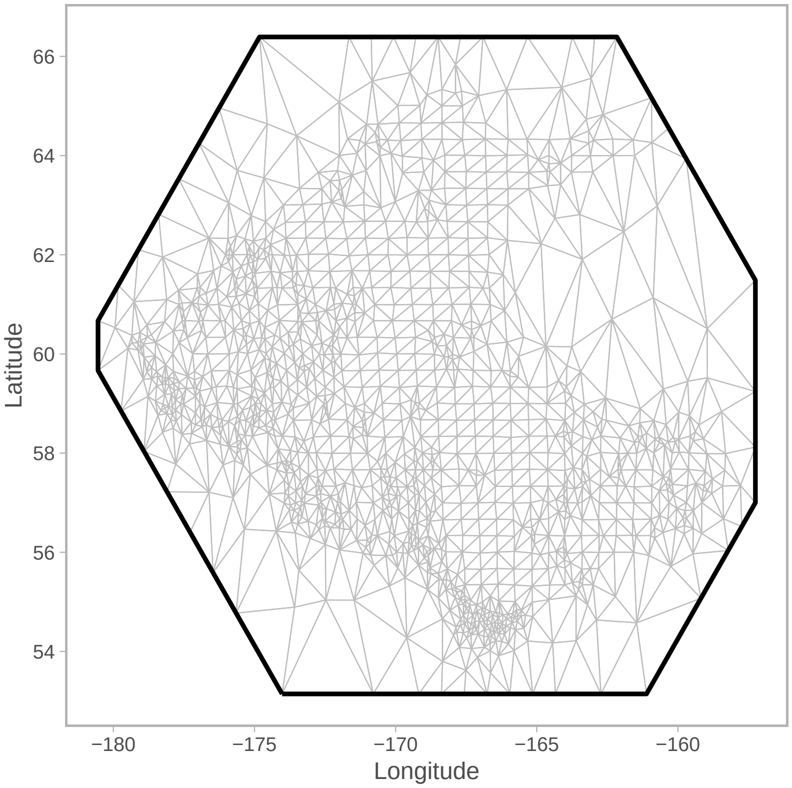


Figure S4: SPDE mesh for the eastern Bering sea case study.

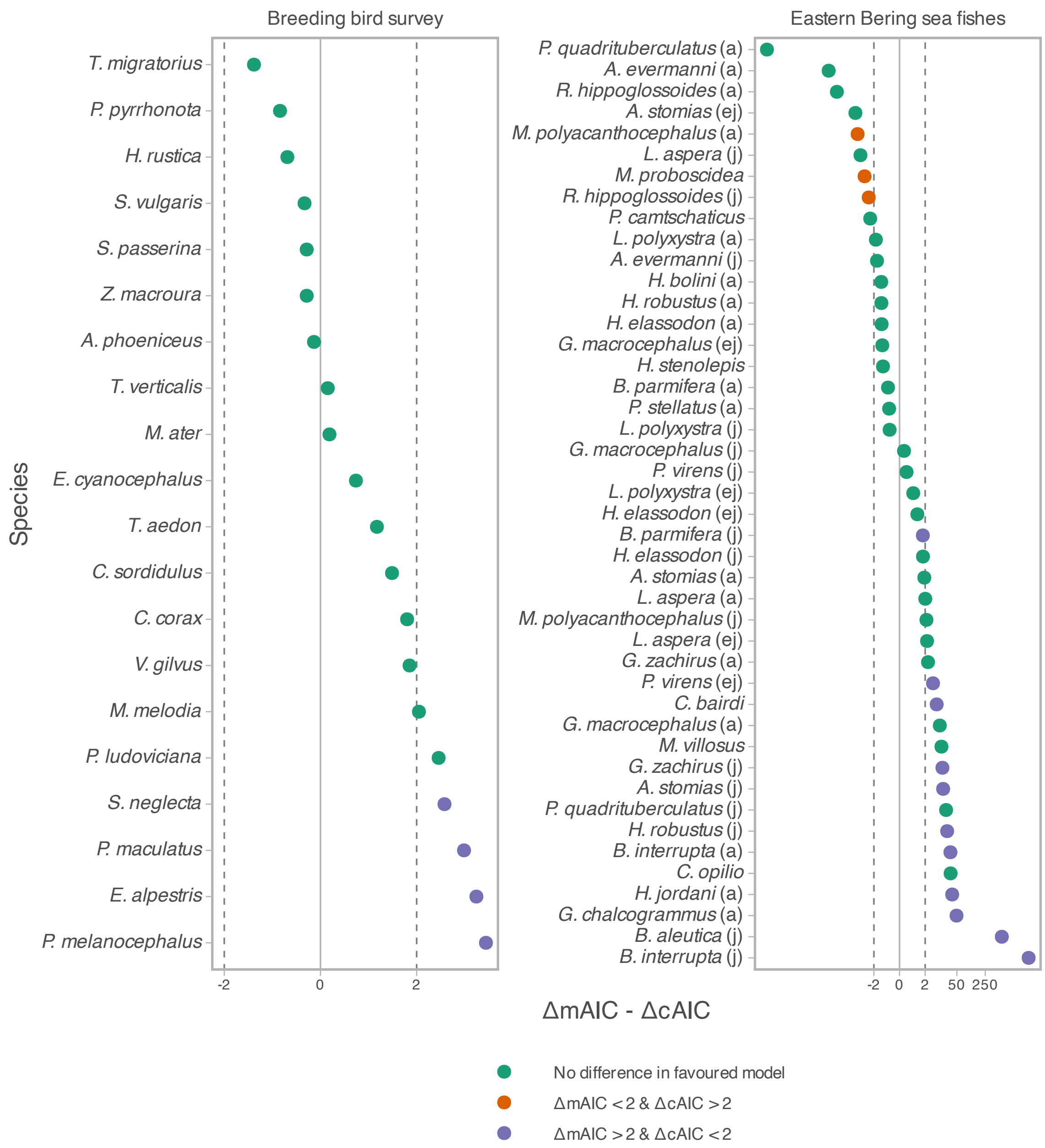


Figure S5: Difference between ΔmAIC and ΔcAIC, where ΔAIC is calculated as the difference between the null model and the diffused model. Point colours indicate whether the two types of AIC lead to consistent selection of the most parsimonious model (diffusion or null). Letters in brackets in the Eastern Bering sea fish case study refers to the life stage (j=juvenile, a=adult, ej = early juvenile). Note the x-axis is fourth-root power transformed.

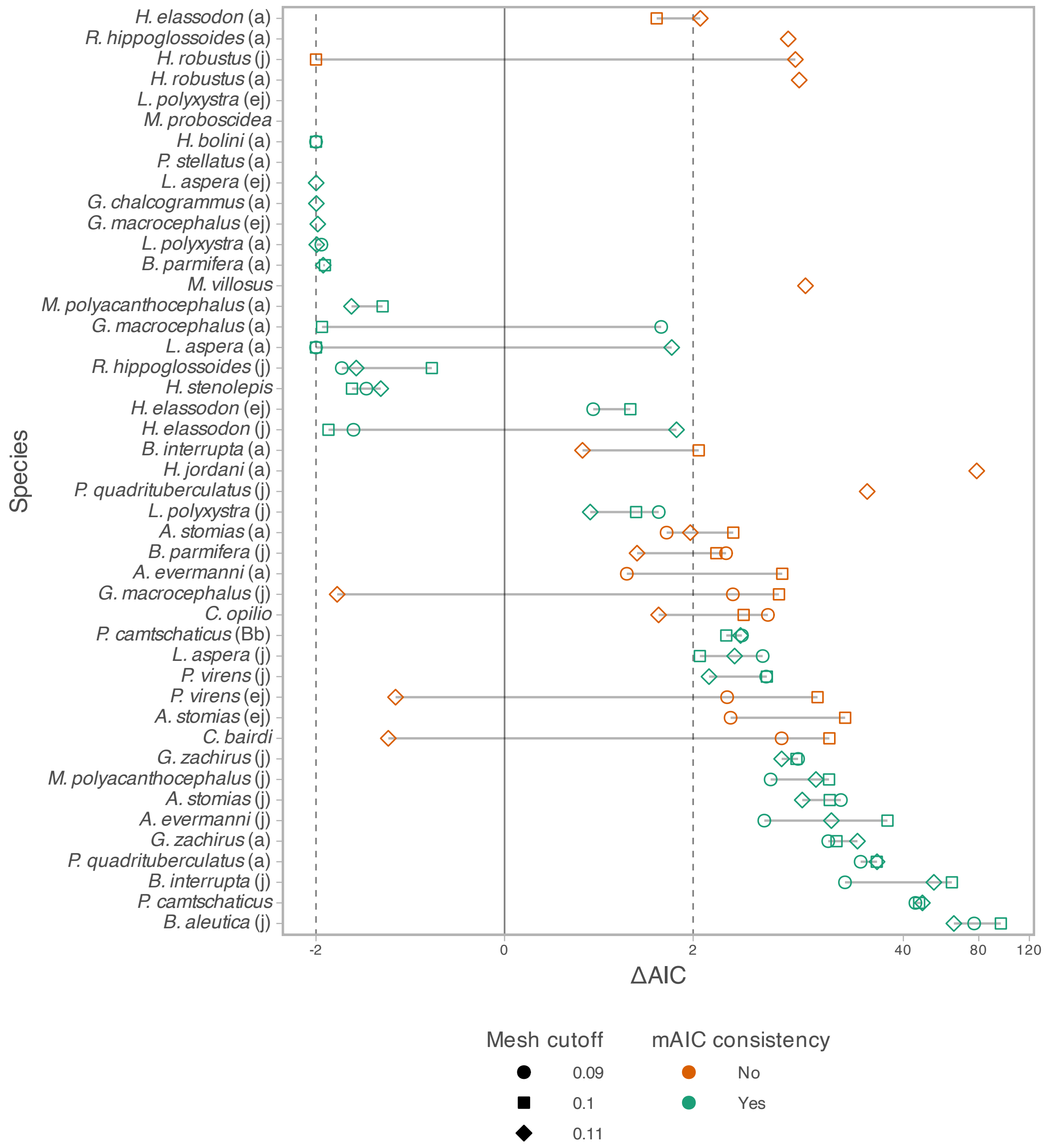


Figure S6: Mesh-sensitivity test showing that for 64% of species-life stage combinations in the Eastern Bering sea fish case study, marginal AIC leads to similar conclusions about which model (null or diffusion) is more parsimonious. Each point represents a single ΔAIC, defined as the difference in marginal AIC between the null and the diffusion model, such that values >2 have support for the diffusion model. Shape indicate the specific mesh resolution (cutoff distance), where low cutoff has has 1475 mesh nodes, cutoff = 1 has 1295, and the high cutoff scenario has 1151 mesh nodes. "mAIC inconsistency=No" refers to the case when any of the 3 mesh-resolution scenarios cross the vertical line at x=2, which indicates that not all models lead to the same conclusion about the diffusion support.

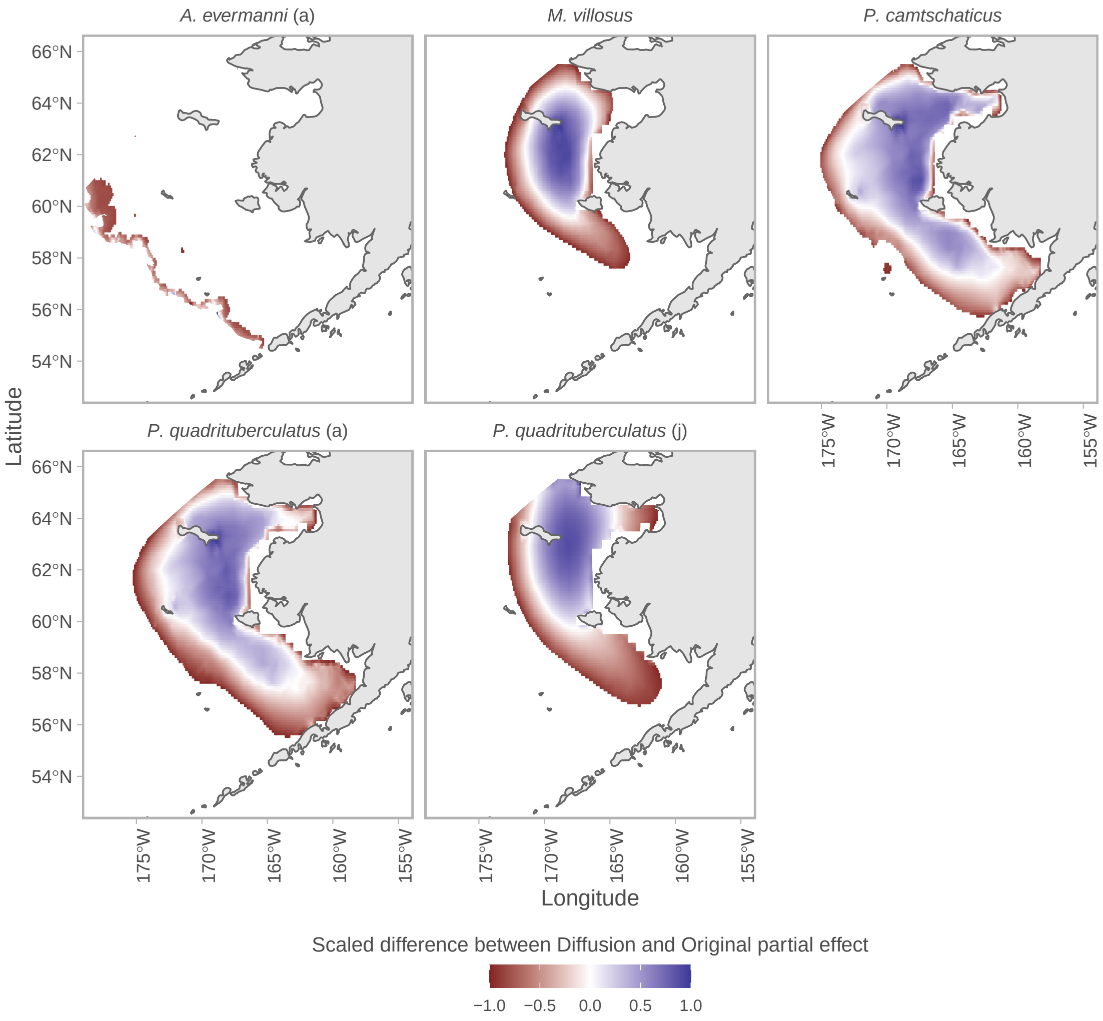


Figure S7: Difference between partial effect of depth from the covariate-diffusion model and the null model for a subset of six fish species from the northeastern Bering Sea case study. For these species, the covariate-diffusion model is supported and the partial predictions suggest they avoid habitats near the continental slope more than inshore areas with a similar depth. Differences in partial predictions are normalized $[-1, 1]$ by species to facilitate comparison among species.

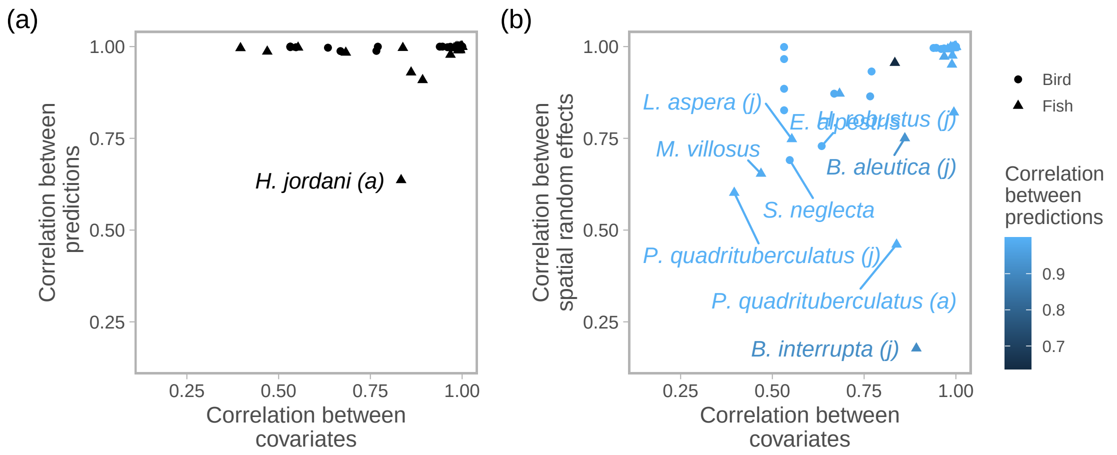


Figure S8: Correlations between predictions from the covariate-diffusion and null model. (a) Predictions from the covariate-diffusion and null model are usually highly correlated regardless of correlation between covariate and diffused-covariate values. (b) There is a positive association between the degree of correlation between covariates (x-axis) and the correlation between spatial random effects (y-axis). Even when the covariate and diffused covariate are not highly correlated, correlations between the predictions can be close to 1. This is presumably due to changes in the spatial random effects (y-axis). For a specific example see Fig. S9.

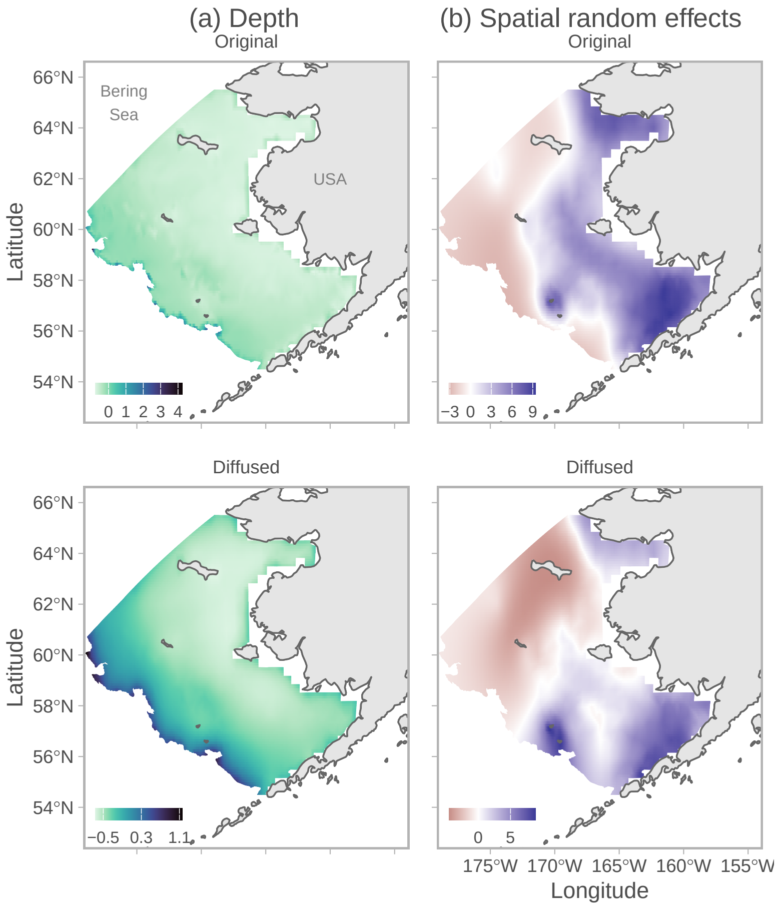


Figure S9: Specific example expanding on Fig. S8 for king crab (*Paralithodes camtschaticus*). Top row is the null model and bottom row is the covariate-diffusion model. (a) Depth covariate and diffused depth covariate. (b) Spatial random effects. (c) Log predicted fish density. The diffused depth covariate is notably different from the raw depth covariate; however, the spatial random effects have also changed resulting in similar predictions.
